# Soil classification predicts differences in prokaryotic communities across a range of geographically distant soils once pH is accounted for

**DOI:** 10.1101/110726

**Authors:** Rachel Kaminsky, Blandine Trouche, Sergio E. Morales

## Abstract

Agricultural land is typically managed based on visible plant life at the expense of the belowground majority. However, microorganisms mediate processes sustaining plant life and the soil environment. To understand the role of microbes we first must understand what controls soil microbial community assembly. We assessed the distribution and composition of prokaryotic communities from soils representing four geographic regions on the South Island of New Zealand. These soils are under three different uses (dairy, sheep and beef, and high country farming) and are representative of major soil classification groups (brown, pallic, gley and recent). We hypothesized that pH would account for major community patterns based on 16S profiles, but that land use and location would be secondary modifiers. Community diversity and structure was linked to pH, coinciding with land use. Soil classification correlated with microbial community structure and evenness, but not richness in high country and sheep and beef communities. The impact of land use and pH remained significant at the regional scale, but soil classification provided support for community variability not explained by either of those factors. These results suggest that several edaphic properties must be examined at multiple spatial scales to robustly examine soil prokaryotic communities.

## Introduction

Sustained population growth has placed a major strain on food production, forcing the development of intensive land use practices that maximize yields^1^. This includes use of heavy machinery and extensive applications of chemical amendments such as fertilizers and herbicides. This intensification of agricultural production has drastically altered soil conditions, causing physicochemical changes (e.g. compaction, decreased organic matter and erosion)^2, 3, 4^ that have led to well-documented losses in biodiversity, including that of belowground microbial communities^5, 6, 7^. Microbes are known to be important to maintaining ecosystem processes^8, 9^. As a result, understanding the consequences of these anthropogenic changes is essential for sustained soil health.

Microorganisms are keystone species that contribute to soil health through bioremediation of contaminants^10, 11,12^ and regulation of nutrient cycling^13, 14,15^. Despite this, the factors that control their distribution and composition are highly contested. Many studies have shown that land use changes influence belowground communities^16, 17, 18^, while pH is a consistent and dominant driver of microbial assemblages on a continental scale and across a range of environments ^19,20,21,22^. However, other edaphic factors like C:N ratio^23^ and soil texture^24,25^ can affect microbial communities. The confounding effects of specific soil factors draws attention to a major gap in prediction and interpretation of microbial community responses to land use change.

Despite the vast number of studies linking individual environmental factors to changes in microbial community structure, the mechanisms underlying these relationships have not been resolved. For example, though there is a widely reported relationship between pH and microbial community structure, it is currently not clear whether pH itself is the most important factor, or if individual chemical and physical factors that contribute to pH are driving this variation^19^. Additionally, many studies concerning land use change focus on a single practice at a particular site^24, 26, 27, 28^. While such analyses provide insight into small-scale microbial community responses to land use intensification, information regarding the comparative responses of communities at multiple scales and across land use types is limited. Moreover, while microbial ecologists seek to capture any and all drivers of belowground communities, it is nearly impossible to measure all environmental factors in a given soil. Most studies evaluate physical factors in terms of soil texture, which is limited in its representation of the complexity of soil. Soil classification provides a more complete description of soils that takes into account the parent material, particle size and permeability, as well as major chemical traits^29^. This parameter also relates soil profiles to climactic and physicochemical features such as weathering, leaching, soil moisture, metal oxides and clay mineral content^30^ and might provide additional resolution for characterizing prokaryotic communities.

To this end, our study used 16S rRNA gene profiles to investigate prokaryotic community composition and distribution in soils on both landscape and regional scales. We examined soils across a series of sites comprising three land use types and four geographic regions. We assess the relationship between prokaryotic communities in these soils with several abiotic factors including pH, land use and soil classification. We hypothesized that prokaryotic community structure would be primarily correlated to pH, while land use would have a secondary relationship with community structure. Furthermore, we hypothesized that soil classification—evaluated at the soil order and subgroup levels—would account for much of the variation in prokaryotic communities not described by either land use or pH. Finally, we sought to understand how individual taxonomic groups responded to these factors.

## Results

### Soil Characteristics

We sampled soils under three land uses: dairy, sheep and beef, and high country. These uses differ in stock type as indicated by their names, but also in their management intensity (i.e. low country = highly managed soils with high stocking rates) as well as location (high country agriculture is carried out on high altitude pastures). Soil physicochemical characteristics varied across land uses, soil order and soil subgroup (Table S1). The sampled soils represented a range of pH values (5.1-6.3). High country soils had, on average, 1.08-fold lower pH than dairy and sheep and beef soils, which were similar in this respect. Soil classification varied within land uses, but most soils are classified within the brown and pallic soil orders, with a few dairy soils representing the recent and gley orders.

### Prokaryotic community structure varies with pH and land use

We examined prokaryotic communities from sites representing three land uses and four geographic regions. A total of 115,445 OTUs (at 97% sequence similarity) were detected within 72 samples representing 24 sites. OTUs per sample ranged between 2,414 and 3,641. Prokaryotic alpha diversity was estimated across all samples and correlations with soil parameters were determined using linear regressions. Richness was correlated with land use (Figure 1A) (Kruskal-Wallis chi-squared = 11.3, *p* < 0.004), with increasing richness from high-country sites to sheep and beef sites. This trend was related to pH (Figure S1A) (regression R^2^ = 0.23, *p* < 0.001) with richness increasing as pH became more neutral. Trends for the Shannon diversity index were similar to those observed for richness with diversity being correlated to both land use (Kruskal-Wallis chi-squared = 26.1*, p* < 0.001) and pH (Figure S1B) (regression R^2^ = 0.48, *p* < 0.001). The remaining chemical data measured in this study (Table S1) did not account for as much variability as pH and land use.

**Figure 1.**
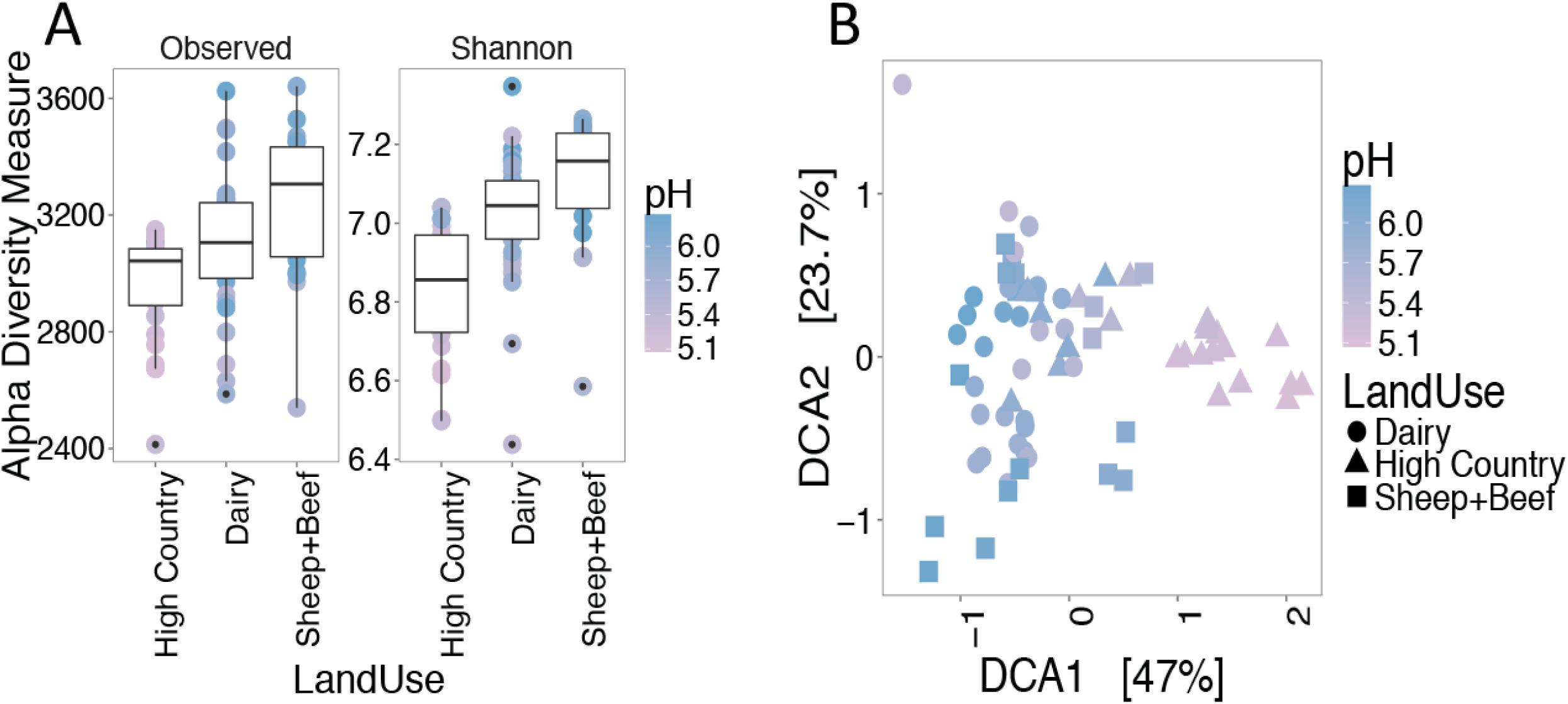
Relationships between bacterial communities under different land uses and pH. Changes in Alpha (Richness and Shannon Diversity) (A) and Beta (Detrended correspondence analysis based on Bray-Curtis dissimilarity) diversity metrics in response to land use and pH (B).

Detrended correspondence analysis (DCA) confirmed trends observed using alpha diversity, with both land use and pH linked to clustering of samples (Figure 1B). Samples from across the three land uses formed a gradient indicating that differences in prokaryotic communities were primarily correlated with changes in pH (Mantel R^2^ = 0.63, *p* < 0.001). While three land uses are included in the study, analysis of similarity (ANOSIM) testing indicated only two major categories: high and low country soils (sheep and beef, and dairy) (Figure S2A, B) (ANOSIM R^2^ = 0.52, *p* < 0.001). Hierarchical clustering of Bray-Curtis distances (Figure S3) confirms the strength of high country and low country environments in explaining the variance in prokaryotic communities (70% confidence). However, sub-clusters representing individual replicates from a site within the high/low country split are better supported using these methods (95% confidence), suggesting unaccounted for factors that are linked to changes in community structure.

### Variation in community composition within land uses is explained by the underlying soil classification

To assess relationships between soil properties and community variation, and observed clustering of samples, within the three land uses data was subset by land use and analyzed independently. Major differences in community structure within the same land use were correlated with soil order, while soil subgroup resolved only a few clusters (Figure 2). Soil subgroup has a significant effect on both the observed species count (Kruskal-Wallis chi-squared = 32.4, *p* < 0.006) and the Shannon diversity index (Kruskal-Wallis chi-squared = 50.6, *p* < 0.001) (Figure 2A). Interestingly, samples grouped based on soil order (Figure 2B) do not have significantly different richness values (*p* > 0.05). However, soil order does correlate weakly with Shannon diversity (Kruskal-Wallis chi-squared = 8.2, *p* < 0.05).

**Figure 2.**
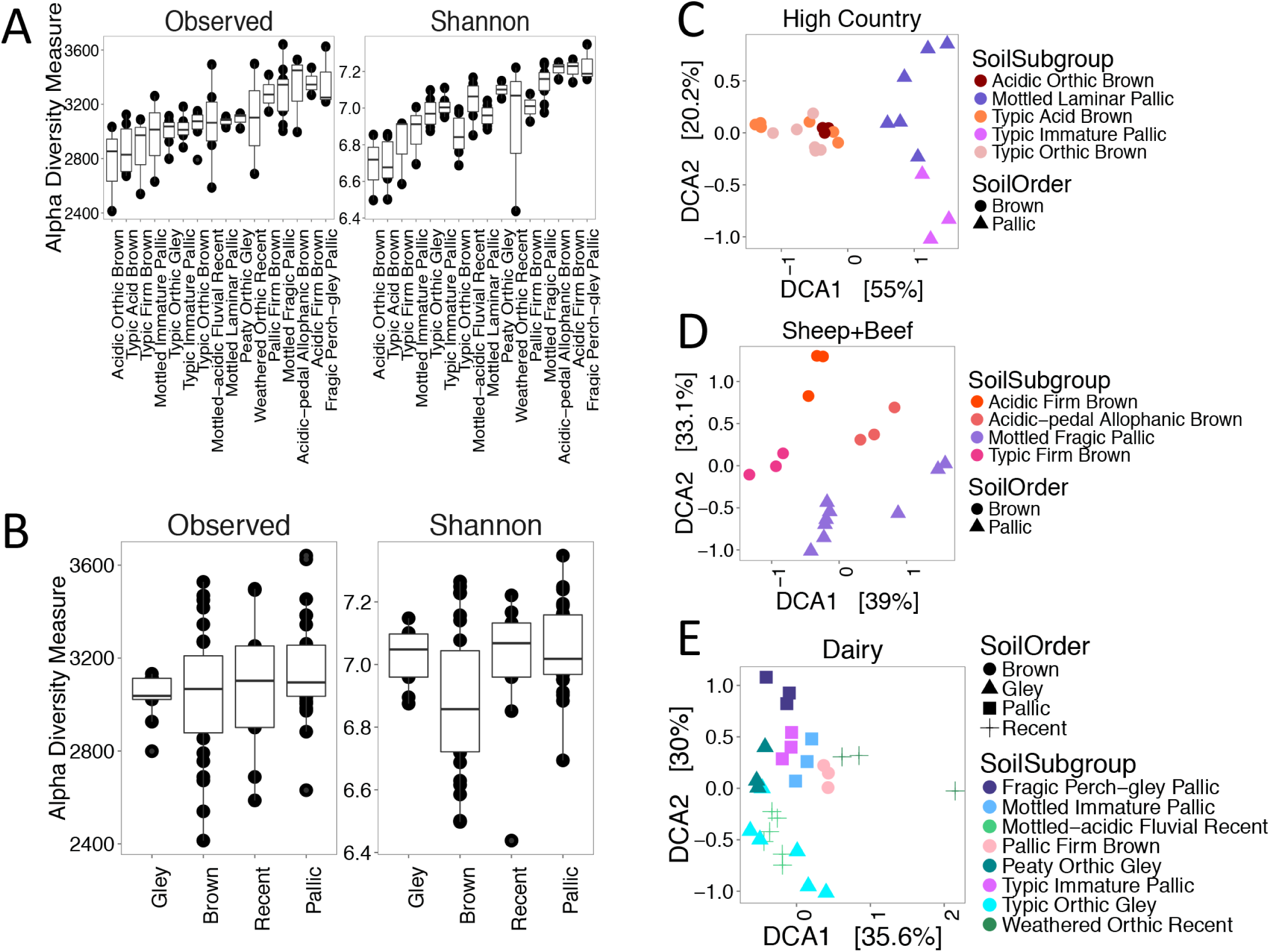
Soil classification predicts prokaryotic community structuring within each land use. Comparison of diversity metrics for each soil subgroup (A) and (B) soil order. High country (C), Sheep and Beef (D), and dairy (E) soil communities evaluated using DCA ordination based on Bray-Curtis dissimilarity with color representing soil subgroup and shape representing soil order.

DCA reveals that prokaryotic communities form distinct clusters based on soil order (Figure 2C, D), though all land use sub-communities have statistically significant relationships with both soil subgroup and soil order (ANOSIM *p* < 0.001). Soil order has a slightly stronger correlation with high country soils (R^2^ = 0.91) (Figure S4A), while sheep and beef communities (R^2^ = 0.58) (Figure S4C) have a slightly stronger relationship with soil subgroup. Hierarchical clustering confirms these results, where high country communities form two clusters (Figure S5), and sheep and beef communities form two (Figure S6). On the other hand, dairy communities do not separate according to soil classification, despite significant correlations with soil order and subgroup (R^2^ = 0.30, 0.67) (Figure S7). These communities remain stable across a wide geographic range, forming one large cluster indicating that an unknown factor reduces variation in dairy soils.

### Influences of pH and land use are stable across multiple spatial scales, but soil classification provides additional support

To determine the impact of geographic scale on observed patterns (based on pH, land use and soil classification), we individually examined the communities from the four geographic regions (Figure S8 and S9). Prokaryotic community changes within regions confirm that pH and land use are the most significant predictors of community structure at multiple scales, while soil classification accounts for the remaining variation (Figure S10-13, Table S2).). Interestingly, land use has the most significant relationships with regional communities where pH was the most significant variable at the multi-region scale.

### Prokaryotic indicators of pH, land use and soil order

Prokaryotic taxa (OTUs) significantly correlated (*p* <0.001) to changes in pH, land use, or soil order were identified using Spearman’s correlations, the Wald test or the Kruskal-Wallis test respectively. The taxa were then mapped onto cladograms (Figure 3; taxa with correlations are provided in Supplementary Table S3).

**Figure 3.**
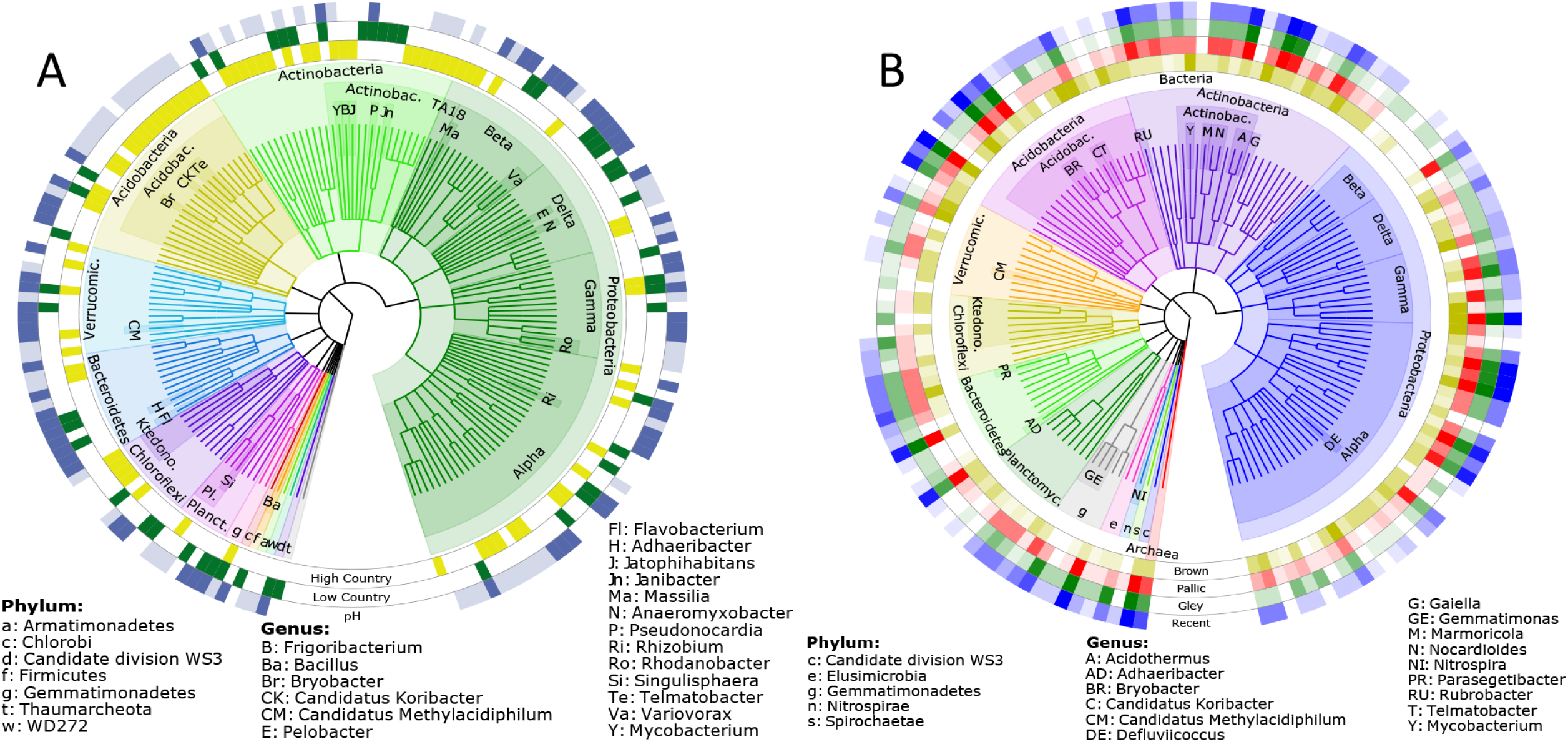
Cladograms showing relationships between key taxa and edaphic properties. (A) OTUs (97% sequence similarity) significantly correlated with high or low country soils and are strongly correlated with changes in pH. Significance for land use preference was determined using the Ward method with a Z lower-limit of 6 and a *p*- value of <0.001. Correlation with pH was determined by a Spearman’s correlation with a Rho lower-limit of 0.5/-0.5 and a *p*-value of <0.001. Light blue indicates a negative correlation with pH, and dark blue is positive (B) OTUs significantly correlated with specific soil orders. Significance was determined using the Kruskal-Wallis test with a chi-squared lower-limit of 27 and a *p*-value of <0.001. Brown soils are indicated by yellow, pallic by red, gley by green and recent by blue. A gradient of 8 shades for each color was generated to indicate abundance, where white indicates an abundance of 0 and the darkest shades indicate an abundance >100.

**Figure 4.**
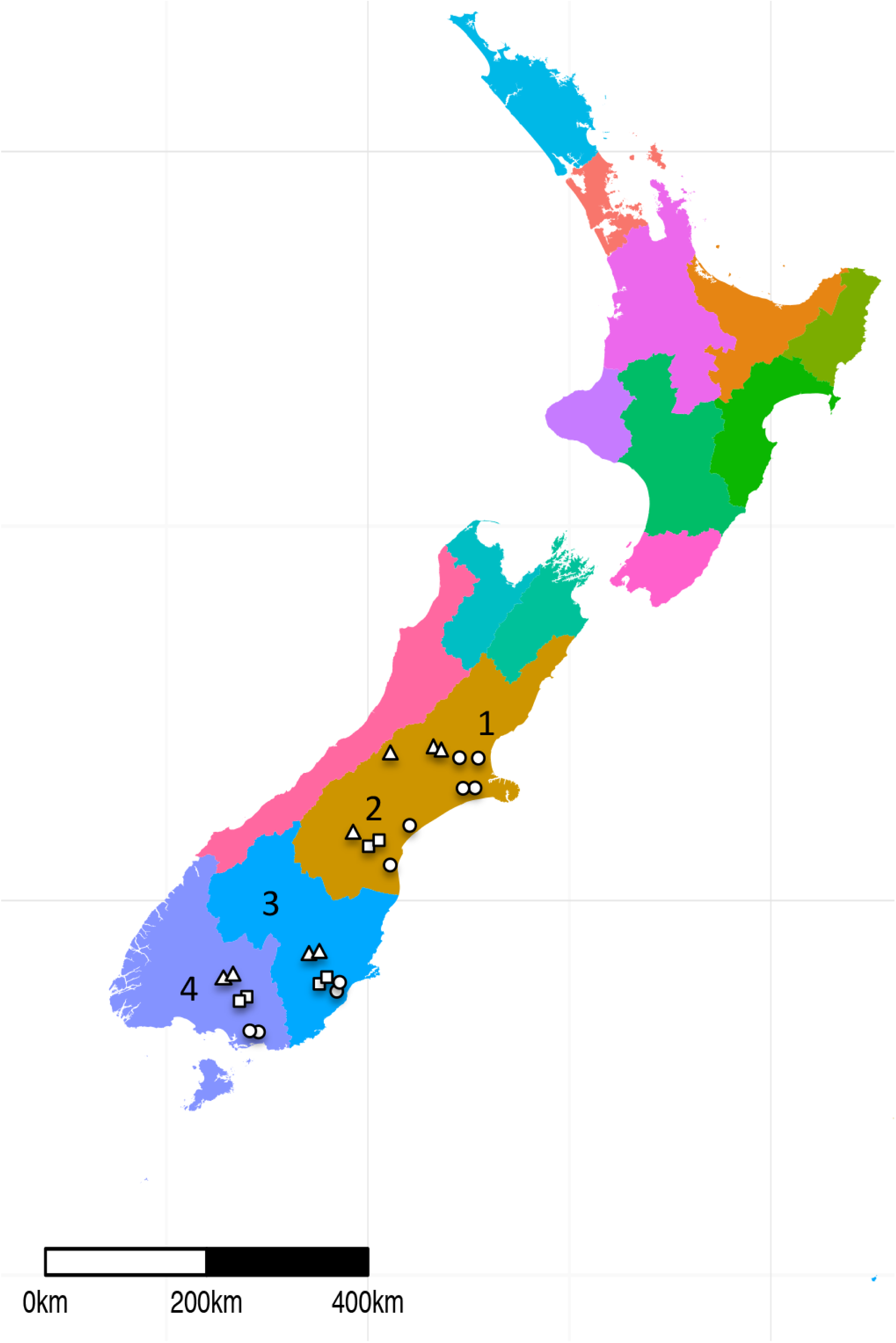
Map of sampling sites throughout the South Island of New Zealand. High country, dairy, and sheep and beef sites are indicated by triangles, circles and squares, respectively. 1-North Canterbury, 2-South Canterbury, 3-Otago, 4-Southland. The map was generated using shapefiles from GADM (v. 2.8, https://www.gadm.org) in RStudio (v. 0.99.903, https://www.rstudio.com/) using ggplot2 (v. 2.1.0), sp (v. 1.2-3), raster (v. 2.5-8), rgdal (v. 1.2-4) and ggsn (v. 0.3.1).

Overall, we found 678 OTUs (0.6% of total OTUs) that were correlated with one or more edaphic properties. 34% of these OTUs correlated with pH, 27% correlated with land use and 40% correlated with soil order. The most represented phyla were the *Proteobacteria* (31% of significant OTUs), *Acidobacteria* (22%), *Actinobacteria* (17%), *Bacteroidetes* (6%) and *Planctomycetes* (5%). A consistent response to specific edaphic properties was not observed at the phylum level.

At the genus level, there was significant overlap between OTU’s identified based on soils classification, pH and land use. Generally, high pH, low country soils, pallic, gley and recent soils shared correlated OTUs (e.g. *Adhaeribacter* and *Revranella)* while low pH, high country soils and brown soils had significantly correlated OTUs in common (e.g. *Bryobacter, Acidothermus, Koribacter, Telmatobacter, Mycobacterium* and *Candidatus Methylacidiphilum).*

However, the relative abundances of several genera correlated with only one edaphic property. *Anaeromyxobacter, Singulisphaera* and *Rhodanobacter* had positive correlations with pH, while *Rhizobium, Variovorax* and *Flavobacterium* were negatively correlated to pH. High country soils were correlated with *Frigoribacterium, Jatrophihabitans* and *Massilia,* while low country soils had correlations with *Janibacter, Pseudonocardia* and *Pelobacter*. Lastly, *Rubrobacter, Defluviicoccus* and *Parasegetibacter* were most strongly correlated with brown soils while *Marmoricola, Nocardiodes* and *Gemmatimonas* had significant correlations with the other three soil orders.

## Discussion

Results revealed that: prokaryotic assemblages differed significantly between land uses and across a pH gradient, however much of the variation within land uses and regions was better accounted for by soil order. Additionally, taxonomic profiles revealed that while overlap exists between OTUs identified as being correlated with pH, land use and soil classification, each parameter identified specific populations not correlated with either of the remaining two.

The studied soils harbored distinct prokaryotic communities, revealing consistent impacts of pH and, to a lesser extent, land use across spatial scales. Our results also confirm the notion that acidic soils support a smaller breadth of diversity. These results are in agreement with many previous studies that have established the role of pH and land use on prokaryotic communities^19, 20, 21^. It has been previously suggested that soil texture is an important predictor of prokaryotic community structure^24, 31, 32^. To build on this relationship, we evaluated the potential link between soil classification (soil order and subgroup) and prokaryotic communities. This allowed us to investigate the extent to which agricultural intensification impacts the relationship between inherent soil properties, like soil texture, and prokaryotic communities. The rationale was that soil classification provides a more thorough representation of the soils’ physical and chemical factors including those not measured (e.g. metal oxides), as well as the geological origins of the soils.

We observed strong relationships between soil classification and prokaryotic community diversity and structure. Brown soils had the lowest diversity, while pallic soils had the highest. The low pH values of the sampled brown soils, combined with the wet climate where some of the brown soils are commonly found^30^, results in low nutrient levels compared to other NZ soils leading to conditions that select for a less diverse community of microbes. In contrast, pallic soils have higher pH values and are only weakly leached, retaining more nutrients allowing for a more diverse community. While richness levels between the two soils were comparable, Shannon diversity differed, indicating changes in evenness. As exemplified by the evenly high levels of iron oxides in brown soils, depleting nutrient stocks and low pH lead to uniform conditions favoring a smaller subset of taxa as shown in our study.

The analysis of sub-communities within each of the four regions suggests that both land use and soil classification have strong relationships with prokaryotic communities. Southland soils had the strongest relationship with land use, but soil order resolved some differences between clusters along the second axis, where communities from a recent soil clustered away from the brown soils. Recent soils are unique in that they are weakly developed, meaning the soil has fewer horizons than the moderately or well-developed soils comprising the other soil orders in this study^33^. Prokaryotic communities from Otago soils were most strongly correlated with soil subgroup. This is especially interesting, as in this region, one of the low country sites grouped with the high country soils on the first DCA axis, but formed their own cluster on the second axis. This cluster happens to contain communities from the only brown soils in this particular region, providing further evidence for soil order as a strong predictor of prokaryotic community structure. In Otago, the two pallic soils clustered quite distantly from one another, explained by the distinction in soil structure between laminar and fragic pallics; laminar soils have layers of clay in the subsoil, while fragic soils are brittle, hard and contain a compacted pan in the subsoil^33^.

Our finding that prokaryotic communities within land uses and regions correlated with soil order indicates that soil classification is a good predictor for prokaryotic communities that are geographically distant from one another. However, we found that dairy communities do not separate clearly based on soil classification. It is possible that the high stocking rates that are characteristic of dairy farms^34, 35^ cause heightened deposition of manure and urine, creating a new soil layer that is fundamentally disconnected from the parent material. It has been shown previously that dairying does impact soil ecosystems in ways that high country, and sheep and beef management does not. For example, Barkle and colleagues observed that application of dairy farm effluent (a mixture of water, urine and manure) onto pasture leads to the accumulation of nutrients and increased prokaryotic biomass^36^. Haynes and colleagues found similar results in camp areas (where livestock tends to congregate) when compared to non-camp soils, which provides further insight into the discrepancy in stocking rate as it affects prokaryotic communities^37^. As a result, the inherent properties expected for soils subjected to dairy management wouldn’t have a relationship with prokaryotic communities. This also gives insight into pH, since soil orders differ in this regard. While it is well established that soil pH is linked to prokaryotic communities on a continental scale, the factors that contribute to pH changes are unresolved ^19^. We can hypothesize that the pH of sheep and beef, and high country soils is connected to inherent soil properties, represented by soil classification, while the pH of dairy soils has been modified by increased agricultural intensification, impacting prokaryotic communities accordingly. Furthermore, while we can be confident in the predictive power of soil order for other land uses, there is less resolution when using soil subgroup. Current methods (charting latitude and longitude onto LRIS soil maps) may not be precise enough to accurately classify soils at this level.

While we have established that pH, land use and soil order are good predictors of prokaryotic community structures, little is known about the mechanisms that account for these relationships. It is possible that pH, land use and soil order serve as integrative variables for multiple chemical and physical characteristics that individually impact prokaryotic communities. Our results suggest that land use, pH and soil order each exert direct pressure on certain prokaryotic taxa, but also contain some overlap in their taxonomic profiles, indicating that they may also integrate some of the same soil properties.

Members of both *Firmicutes (Bacillus)* and *Thaumarchaeota* (uncultured representative) are significantly represented in low country soils, but not at high pH levels. This is interesting, as many members of these phyla are thought to thrive at high pH levels^38, 39^, suggesting that the members detected here have different life strategies that are selected for by land use. Additionally, DCA plotting showed that high country soils are strongly correlated with low pH, which is supported by their shared relationship with several *Acidobacteria* groups. However, there were several members from the *Proteobacteria* (e.g. *Massilia), Actinobacteria* (e.g. *Frigoribacterium),* and *Chloroflexi* (e.g. *Ktedonobacter*) that were significantly represented in high country soils but not at low pH levels. Little is known about the ecophysiology of many of these genera. However, *Massilia* are copiotrophs, and are sensitive to nutrient availability. It is established that high country rangelands are subjected to less rigorous management regimes compared to their low country counterparts^40^. This management strategy may give rise to a nutrient profile that is preferable for the maintenance of *Massilia* populations^41^. Selection by land use is further evidenced by the strong correlation between high country soils and the verrucomicrobial phylotype Da101 and, contrastingly, a positive correlation with pH. As high country soils tend to have lower pH values, and *Verrucomicrobia* are thought to persist in low-nutrient environments^42, 43^, it can be inferred that the stable nutrient status of high country soils explains the abundance of this phylotype rather than pH. Other taxa, like *Gaiella* (originally isolated from an aquifer^44^) and *Nitrospira,* which are normally found in wet environments^45^, were most significantly correlated with gley soils. These soils are known to have high water tables^30^, which would likely provide preferable conditions for these microbes to thrive.

Our results confirm soil pH is the strongest predictor of community structure, diversity and composition across multiple spatial scales, but we also show strong relationships with land use and soil order. We propose that soil order may serve as an integrative factor that accounts for physical and chemical properties and can be used when direct assessment of specific edaphic factors is not possible. Further, the identification of specific OTUs correlated to more than one factor suggests that spurious correlations are highly likely and other factors besides pH might better explain observed patterns.

## Materials and methods

### Soil sampling

A total of 24 field sites across four regions on the south island of New Zealand were sampled in this study (Figure 1). Sites were chosen to represent: the three main land uses in New Zealand agriculture (dairy, sheep and beef, high country farming), a wide range of edaphic parameters (Table S1), and four major regions of New Zealand (North Canterbury, South Canterbury, Otago, Southland). Samples were collected at the beginning of the growing season, between May 5 and May 16, 2014. Sites were delineated in the field by twelve replicate plots (1m^2^ each) within a gridded area enclosed by a 6.5 by 5 m fence. Biological replicates from each site were collected by sampling three random plots for a total of 72 samples in the study (24 sites × 3 plots at each site). Each sample comprised a composite of four cores (7.5 cm depth and 2.5 cm diameter) that were taken 0.4 m apart diagonally across the 1m^2^ plot. Cores were screened prior to compositing to remove roots, worms and rocks. Samples were kept on ice while in the field and stored at -20 degrees until returning to the lab for final storage at -80 degrees.

Chemical analyses were performed by R.J. Hill Laboratories (Hamilton, NZ). For soil pH determination, a 1:2 soil: water slurry was prepared followed by potentiometric titration (CITE). Data for soil physical properties were obtained from the New Zealand Land Resource Information Systems Portal (https://lris.scinfo.org.nz/).

### DNA extraction and sequencing

Genomic DNA was extracted from 0.25 g of soil using the Mo Bio PowerSoil-htp 96-well soil DNA isolation kit (Carlsbad, CA, USA) according to the manufacturer’s instructions, but with a modification at the lysing step. Samples were placed on a Geno/Grinder homogenizer (SPEX Sample Prep, LLC, Metuchen, NJ, USA) for two rounds of fifteen seconds at 1750 strokes/minute. One extraction was performed on each sample. DNA concentration and purity was determined using a Nanodrop 1000 Spectrophotometer (Thermo Scientific, Wilmington, DE, USA). Absorbance was observed at 230, 260, 280 and 320 nm.

The V4 region of the 16S rRNA gene was amplified using the universal primer pair 515F (5'-NNNNNNNNGTGTGCCAGCMGCCGCGGTAA-3') and 806R (5'-GGACTACHVGGGTWTCTAAT-3') following the Earth Microbiome Project barcoded conditions^46^. Each sample was given a barcode sequence on the 5’ end of the forward primer for multiplexed sequencing and loaded onto a single Illumina MiSeq 2 × 151 bp run (Illumina, Inc., CA, USA). Sequences were deposited at the Sequence Read Archive (NCBI) with the accession numbers: 5902515-5902586 under the BioProject ID: PRJNA348131.

### Sequence processing

All sequences were initially processed using a QIIME 1.9.0 open-reference OTU-picking workflow^47^. In brief, raw sequences were first demultiplexed. Forward sequences were then clustered into OTUs (97% similarity) against the SILVA database release 119^48^ using UCLUST^49^. Reads that failed to hit the reference database were clustered *de novo.* Taxonomy assignments were determined using BLAST^50^ with a maximum e-value of 0.001 against the SILVA database. The resulting OTU table was then subsampled to an even depth of 12,000 sequences per sample ten times followed by merging of the resulting ten OTU tables to reduce biases that arise from unequal library sizes. All data was then exported as a biom file.

### Statistical analyses

Sample counts were transformed by dividing the individual OTU abundances by the number of rarefactions (10) followed by rounding prior to downstream analysis using the phyloseq package^51^ in R^52, 53^. Diversity estimates were determined using observed richness and the Shannon index, as calculated and plotted in phyloseq and ggplot2^54^ Regression analyses and Kruskal-Wallis tests were performed in R to assess the relationships between environmental variables and richness and diversity. Prokaryotic community differences were represented on a two-dimensional ordination plot using Detrended Correspondence Analysis (DCA) with the Bray-Curtis distance between samples in phyloseq and ggplot2. Analysis of Similarity (ANOSIM) was used to quantify the relationships between significant differences in community structure and categorical variables (land use and soil classification) within the vegan package^55^. The Mantel test was performed in vegan with 999 permutations to assess relationships between continuous variables (pH) and community structure. To identify consistent clustering patterns in the data, hierarchical clustering was performed in the pvclust package^56^ using Ward’s method and Bray-Curtis distances. To examine significant differences in the abundance and distribution of taxa between land uses, the data were transformed to relative abundance in phyloseq. The Wald chi-squared test was applied to the data using the DESeq2 package^57^. Spearman’s rank correlations were used to test differences in taxa distributions along the pH gradient. The Kruskal-Wallis test was used to observe differences in OTU abundances of significance between the soil orders, and was performed in QIIME. Cladograms were generated in GraPhlAn^58^. Mapping was done using GADM^59^ in RStudio with packages: ggplot2, sp^60, 61^, raster^62^, rgdal^63^ and ggsn^64^.

## Acknowledgements

We thank Mainland Minerals Ltd. for funding this research and assisting with soil sampling. We also thank Hill Laboratories and Soiltech for providing physicochemical analyses and Dr. Xochitl Morgan for her helpful comments on this manuscript. Rachel Kaminsky was funded through a Callaghan Innovation education fellowship (MMSOU1301).

## Author Contributions

S.E.M. designed the experiment. R.K collected and processed samples. R.K., S.E.M and B.T. analyzed data. All authors were involved in the writing process.

## Additional Information

### Accession codes

Sequences were deposited at the Sequence Read Archive (NCBI) with the accession numbers: 5902515-5902586 under the BioProject ID: PRJNA348131.

